# INPP5E interactome reveals novel connections to growth factor signaling

**DOI:** 10.64898/2026.02.13.705725

**Authors:** Raquel Martin-Morales, Pablo Barbeito, Belén Sierra-Rodero, Lucia Jimenez, Maria Schrøder Holm, Darío Cilleros-Rodríguez, Lena K. Ebert, Matilde Cortabarría, Lotte B. Pedersen, Jose L. Badano, Florencia Irigoín, Bernhard Schermer, Wolfgang Link, Søren T. Christensen, Francesc R. Garcia-Gonzalo

**Affiliations:** Instituto de Investigaciones Biomédicas Sols-Morreale (IIBM), UAM-CSIC, Madrid, Spain; Departamento de Bioquímica, Facultad de Medicina, Universidad Autónoma de Madrid (UAM), Madrid, Spain; CIBER de Enfermedades Raras (CIBERER), Instituto de Salud Carlos III, Madrid, Spain; Department of Biology, University of Copenhagen, Copenhagen, Denmark; Department II of Internal Medicine and Center for Molecular Medicine Cologne, University of Cologne, Faculty of Medicine and University Hospital Cologne, Cologne, Germany; Cologne Excellence Cluster on Cellular Stress Responses in Aging-Associated Diseases (CECAD), University of Cologne, Faculty of Medicine and University Hospital Cologne, Cologne, Germany; Human Molecular Genetics Laboratory, Institut Pasteur de Montevideo, Montevideo, Uruguay; Unidad Académica de Histología y Embriología, Facultad de Medicina, Universidad de la República, Montevideo, Uruguay

**Keywords:** INPP5E, cilia, Joubert syndrome, growth factors, TGFβ signaling, PDGF signaling

## Abstract

Primary cilia are sensory cell membrane protrusions whose malfunction causes diseases known as ciliopathies. Joubert syndrome (JBTS) is a rare recessive ciliopathy causing brain malformations and kidney cysts, among other manifestations. A key player in JBTS is INPP5E, a ciliary phosphoinositide lipid phosphatase. Although INPP5E regulates ciliary growth factor signaling, the molecular mechanisms remain poorly understood. Herein, we show that a constitutively active growth factor receptor (PDGFRα-D842V) stimulates INPP5E tyrosine phosphorylation, as does the SRC tyrosine kinase. INPP5E tyrosine phosphorylation did not affect its enzyme activity but was associated with stronger binding to SH3GL1, an endocytic regulator, and SNX9, a phosphoinositide-binding protein involved, like INPP5E, in growth factor-induced ciliary ectovesicle release. Our INPP5E interactomic studies identified other growth factor signaling regulators, including among others: SIN1 (the phosphoinositide-binding subunit of the mTORC2 complex), STRAP (a TGFβ and PI3K/AKT signaling regulator), GRB2 (a growth factor receptor adaptor) and the JBTS-associated proteins AHI1 and NPHP1. Moreover, INPP5E strongly interacted with the phosphopeptide-binding 14-3-3 proteins, and did so only in presence of serine-85, a phosphorylated INPP5E residue. Lastly, we found that INPP5E differentially regulates two cilium-dependent growth factor signaling pathways in fibroblasts. Thus, while INPP5E downregulated PDGF-induced AKT phosphorylations, it upregulated TGFβ-induced SMAD2 and ERK phosphorylations. Altogether, our work provides important clues on the cilia-dependent actions of growth factors, with implications for ciliopathies like JBTS.

## Introduction

Cilia are finger-like plasma membrane appendages formed by microtubules emanating from the basal body, a membrane-anchored centriole. Many mammalian cell types form cilia, which they use to either displace extracellular fluids (motile cilia) or mediate responses to extracellular cues (primary cilia). Some of these cues include key developmental regulators, like Hedgehog (Hh) and Wnt ligands, and growth factors such as transforming growth factor beta (TGFβ) and platelet-derived growth factor (PDGF). Primary cilia malfunction is linked to cancer, obesity and genetic disorders called ciliopathies [1, 2].

Joubert-Boltshauser syndrome (JBTS), or simply Joubert syndrome (JS), is a rare recessive ciliopathy with a prevalence around 1 in 100,000. JBTS manifestations include brain malformations, motor and cognitive deficits, kidney cysts, polydactyly and retinal degeneration [3]. Genetically, JBTS is caused by mutations in over 40 ciliary genes. Of these, *INPP5E*, encoding a ciliary membrane-localized phosphoinositide 5-phosphatase, is one of the most commonly affected.

The INPP5E protein consists of an N-terminal proline-rich disordered region, the 5-phosphatase catalytic domain, and a C-terminal CaaX box, whose farnesylation contributes to membrane anchoring and ciliary targeting. While mutations in INPP5E’s catalytic domain lead to JBTS or non-syndromic retinal degeneration, mutations in its CaaX box cause MORM syndrome, a different ciliopathy characterized by mental retardation, obesity, retinal degeneration and micropenis [3–6].

Of the 40+ JBTS-causative proteins, about half are needed for INPP5E ciliary accumulation. These include the ciliary membrane protein ARL13B, the farnesyl carrier PDE6D, and multiple components of the transition zone (TZ), the structure that forms the ciliary gate separating the cilium from the rest of the cell. Such TZ components include known INPP5E interactors like AHI1 and NPHP1 [3, 4, 7–12].

Once inside cilia, a major function of INPP5E involves dephosphorylating phosphatidylinositol-4,5-bisphosphate (PI(4,5)P_2_), a phosphoinositide lipid whose ciliary levels must be kept low for proper function of Hh signaling, an important pathway in ciliopathies. Accordingly, INPP5E-knockout (KO) cells display abnormal ciliary accumulations of PI(4,5)P_2_ and the PI(4,5)P_2_-dependent Hh pathway repressors TULP3 and GPR161, resulting in poor Hh responsiveness [13–19].

Several lines of evidence also point to important roles for INPP5E in growth factor signaling. First, INPP5E can also dephosphorylate phosphatidylinositol-3,4,5-trisphosphate (PIP_3_), a key second messenger in growth factor-stimulated PI3K-AKT signaling. Consistently, AKT is overactive in INPP5E-KO kidneys and brain [18, 20–26].

Moreover, INPP5E regulates cilia disassembly and ciliary ectovesicle (EV) release, two growth factor-dependent processes. Cilia disassembly is accelerated in INPP5E-KO cells in a manner dependent on Aurora kinase A (AURKA), a key cilia disassembly regulator with which INPP5E physically and functionally interacts [5, 6, 27]. Ciliary EV release is triggered by growth factors raising ciliary PI(4,5)P_2_, which then promotes F-actin-dependent vesicle scission. High PI(4,5)P_2_ levels in INPP5E-KO cilia make EV release constitutive in these cells [28, 29].

Despite all the evidence connecting INPP5E to growth factor signaling, many questions remain. Herein, we mainly address two: (i) how growth factors regulate INPP5E’s activity and interactions; and (ii) which growth factor pathways depend on INPP5E.

First, we show that a constitutively active PDGF receptor mutant, PDGFRα-D842V, induces strong tyrosine phosphorylation of INPP5E. We then characterized the INPP5E interactome in presence or absence of this D842V mutant. Although most identified interactors showed no D842V-dependence, a few did, most notably SNX9, a phosphoinositide-binding F-actin regulator implicated in ciliary EV release [28]. The strongest INPP5E interactors were the 14-3-3 proteins, phosphopeptide-binding adaptors whose interaction we demonstrate requires INPP5E’s serine-85 (S85), a phosphorylated residue part of a canonical 14-3-3-binding site [30]. INPP5E’s proline-rich region also contains several predicted Src Homology 3 (SH3)-binding motifs, and we accordingly find that INPP5E interacts with multiple SH3 domain-containing proteins, including SNX9, GRB2, SH3GL1, NPHP1 and AHI1 [31]. Other INPP5E interactors we validated included SIN1 (the mTORC2 complex’s phosphoinositide-binding subunit) and STRAP (a phosphoinositide and TGFβ signaling regulator) [32–35]. Consistently, we show that INPP5E regulates TGFβ and PDGF signaling pathways, both of which are cilium-dependent [2, 36–44].

## Methods

### Antibodies and Reagents

Mouse anti-alpha-tubulin (Proteintech, 66031-1-Ig), mouse anti-EGFP (Proteintech, 66002-1-Ig), rabbit anti-EGFP (Proteintech, 50430-2-AP), mouse anti-Flag (Proteintech, 66008-3-Ig, or Sigma, F3165 or F1804), rabbit anti-Myc (Proteintech, 16286-1-AP) and mouse anti-V5 (Thermofisher, MA5-15253) were used as described [4, 45]. Also used as reported were goat anti-PDGFRα (R&D Systems, AF1062), rabbit anti-phospho-Rb (Cell Signaling, 9308), rabbit anti-phospho-SMAD2 (Cell Signaling, 3108), rabbit anti-SMAD2 (Abcam, ab40855), rabbit anti-phospho-ERK1/2 (Cell Signaling, 9101), rabbit anti-ERK1/2 (Cell Signaling, 9102), mouse anti-DCTN1 (p150glued) (BD Bioscience, 612708) and rabbit anti-phospho-SRC (Y419) (Invitrogen, 44-660G) [40, 41, 43, 46–48].

The following rabbit antibodies were from Proteintech and were used at dilution 1:1000, unless otherwise specified: anti-AHI1 (22045-1-AP), anti-14-3-3ζ (14881-1-AP), anti-USP9X (55054-1-AP), anti-SIN1 (15463-1-AP; WB: 1:500), anti-CAMSAP2 (17880-1-AP), anti-SH3GL1 (27014-1-AP), anti-SNX9 (15721-1-AP), and anti-STRAP (18277-1-AP). Other antibodies included: mouse anti-phosphotyrosine (PY20, Biolegend, 309302, WB: 1:1000), rabbit anti-AKT (Cell Signaling, 9272, WB: 1:1000), rabbit anti-phospho-AKT (S473) (Cell Signaling, 4060, WB: 1:1000), rabbit anti-phospho-AKT (T308) (Cell Signaling, 4056, WB: 1:1000), and mouse anti-beta-actin (Proteintech, 66009-1-Ig, WB: 1:10,000). All secondary antibodies were as previously reported [4, 45].

Other reagents included GFP-Trap_MA beads (Proteintech, gtma), mouse anti-Flag M2 magnetic beads (Sigma, M8823), sodium orthovanadate (Alfa Aesar, J60191) and the following recombinant human growth factors: PDGF-AA (Proteintech, HZ-1215), TGF-β1 (R&D Systems, 240-B), Insulin (Gibco, 12585014), IGF-I (Proteintech, HZ-1322) and EGF (Proteintech, HZ-1326).

### Plasmids

Plasmid encoding V5-INPP5E was obtained using a previously described modified pcDNA6 vector (Seeger-Nukpezah et al. 2012). Flag-INPP5E, EGFP-INPP5E and all the latter’s CLS and deletion mutants are described elsewhere [4]. To make the S85A, S99A, S85A+S99A, Y293F and Y614F mutants of EGFP-INPP5E, we used overlap extension PCR, as before [4]. Also reported are 5HT_7_-EGFP [49], EYFP-SNX9 [28], Flag-EPS15L1(1-225), and Flag-NPHP1 and its mutants [50]. To generate NPHP1-myc plasmid, NPHP1 coding sequence was amplified from Flag-NPHP1 (pcDNA6-Flag-His-NPHP1) and cloned into pcDNA3.1-myc-his(-)C using BamHI-KpnI sites. Plasmids encoding PDGFRα-EGFP (WT and D842V) and PDGFRβ-EGFP in the pcDNA5FRT-EF backbone are also reported (Addgene #66787, 66789 and 66790) [51]. To make PDGFRα-Flag (WT & D842V), the 66787 and 66789 plasmids were opened with KpnI and ligated to pre-annealed primers encoding the Flag tag followed by stop codons in all three reading frames, all flanked by KpnI cohesive ends on both sides. For Flag-AHI1, the BamHI insert from EGFP-AHI1 (Addgene #30494) was subcloned into the pFlag-CMV4 vector [52]. Plasmids encoding chicken c-Src (pcDNA3-c-SRC-WT), EYFP-GRB2 and EYFP-SH3GL1 were gifts from Drs. Jorge Martín-Pérez, José M. Rojas and Takanari Inoue, respectively [28, 48, 53].

### Cell culture, transfections and treatments

HEK293T, hTERT-RPE1, *Inpp5e* ^+/‒^ and *Inpp5e* ^‒/‒^ MEFs, and puromycin-sensitive WT and INPP5E-KO hTERT-RPE1 cells were cultured and transfected as previously reported [4, 13, 45]. For growth factor treatments, MEFs or hTERT-RPE1 cells were serum-starved for 24 hours in basal medium (DMEM for MEFs; DMEM/F12 for RPE1)+0.2% FBS (starvation medium) and thereafter treated in starvation medium with either 50 ng/ml PDGF-AA, 2 ng/ml TGF-β1, 100 nM insulin, 10 nM IGF-1, 50 ng/ml EGF or 15% FBS. Cells were then lysed in RIPA buffer containing Halt phosphatase and protease inhibitor cocktails (Thermofisher) and processed for Western blot analysis. For orthovanadate treatment, final concentrations of 10 μM Na_3_VO_4_ and 3 mM H_2_O_2_ were added to OptiMEM medium, and the pervanadate-containing medium was immediately used to treat cells for 10 min at 37°C before cell lysis as above [54].

### Immunoprecipitation, Western blot and Proteomics

Immunoprecipitations from transfected HEK293T cell lysates were performed with GFP-Trap_MA beads (Proteintech, gtma) or mouse anti-Flag M2 magnetic beads (Sigma, M8823) following previously reported protocols, which also described Western blotting and its quantitation [4, 45]. For the proteomics experiment, HEK293T cells, three 15-cm plates per condition, were transfected with: (i) pFlagCMV4 + PDGFRα-D842V-EGFP, (ii) Flag-INPP5E + PDGFRα-D842V-EGFP, or (iii) Flag-INPP5E + PDGFRα-WT-EGFP. Cells were lysed at 40 hours post-transfection with buffer containing 50 mM Tris-HCl pH 7.5, 150 mM NaCl, 1% Igepal CA-630 (Sigma) and 1X Halt protease and phosphatase inhibitor cocktails (Thermofisher). After overnight immunoprecipitation, the Flag M2 beads were washed thrice in lysis buffer minus inhibitors, eluted in Laemmli buffer, and submitted to the proteomics facility of the Spanish National Center for Biotechnology (CNB-CSIC), where samples underwent SDS-PAGE, tryptic digestion, liquid chromatography and tandem mass spectrometry (LC-ESI-MS/MS) for protein identification, as previously described [45].

### Phosphoinositide phosphatase activity assays

Activity assays were performed as previously described [4]. Briefly, lysates from transfected HEK293T cells were subjected to immunoprecipitation with GFP-Trap_MA beads (Chromotek), after which the EGFP-INPP5E-containing beads were washed and incubated with reaction buffer including diC8-PtdIns(4,5)P2 or diC8-PIP3 as substrates (Echelon Biosciences). After 20 min of reaction at 37 °C, phosphate release was measured in the supernatant using the Malachite Green Assay Kit (Echelon Biosciences).

### Quantitations and statistics

Western blot quantitations were done with Fiji/ImageJ as previously described [4]. To compare growth factor responses across experiments, phosphorylation ratios in each experiment (R=pAKT/AKT; R=pSMAD2/SMAD2; or R=pERK/ERK) were normalized according to:

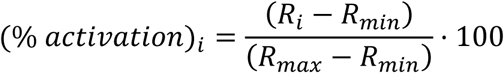

In this equation, R_i_ is the phosphorylation ratio for sample i, while R_max_ and R_min_ are the maximum and minimum ratios, respectively, of that particular experiment. Hence, the above equation assigns 0% to the minimum value, 100% to the maximum, and intermediate values to all other samples in the experiment, reflecting relative phosphorylation levels. In these and other experiments, GraphPad Prism 10 was used for graphs and statistical analyses, whose relevant details are provided in the figure legends.

## Results

### INPP5E undergoes tyrosine phosphorylation

Given INPP5E’s involvement in growth factor-induced cilia disassembly [5, 6, 28], we wondered whether INPP5E could be phosphorylated in presence of PDGFRα-D842V, a cilia disassembly-inducing PDGFRα mutant with constitutive tyrosine kinase activity [51]. To test this, we used immunoblot to detect the phosphotyrosine (pY) associated with EGFP-tagged INPP5E when the latter was immunoprecipitated from HEK293T cell lysates also expressing Flag-tagged PDGFRα-D842V, or its wild type (WT) form as control (Fig. 1a). Strong pY signal at the exact position of the EGFP-INPP5E band was observed with Flag-PDGFRα-D842V, but not with Flag-PDGFRα-WT or empty vector (Fig.1a). Coexpression of wild type SRC tyrosine kinase also caused EGFP-INPP5E to display positive pY signal, albeit less intensely than with D842V (Fig.1a). Consistently, the overexpressed SRC-WT showed positivity for phosphotyrosine-418 (pY418), indicating that a subset of the kinase was in its active conformation (Fig.1a) [48].

**Figure 1.**
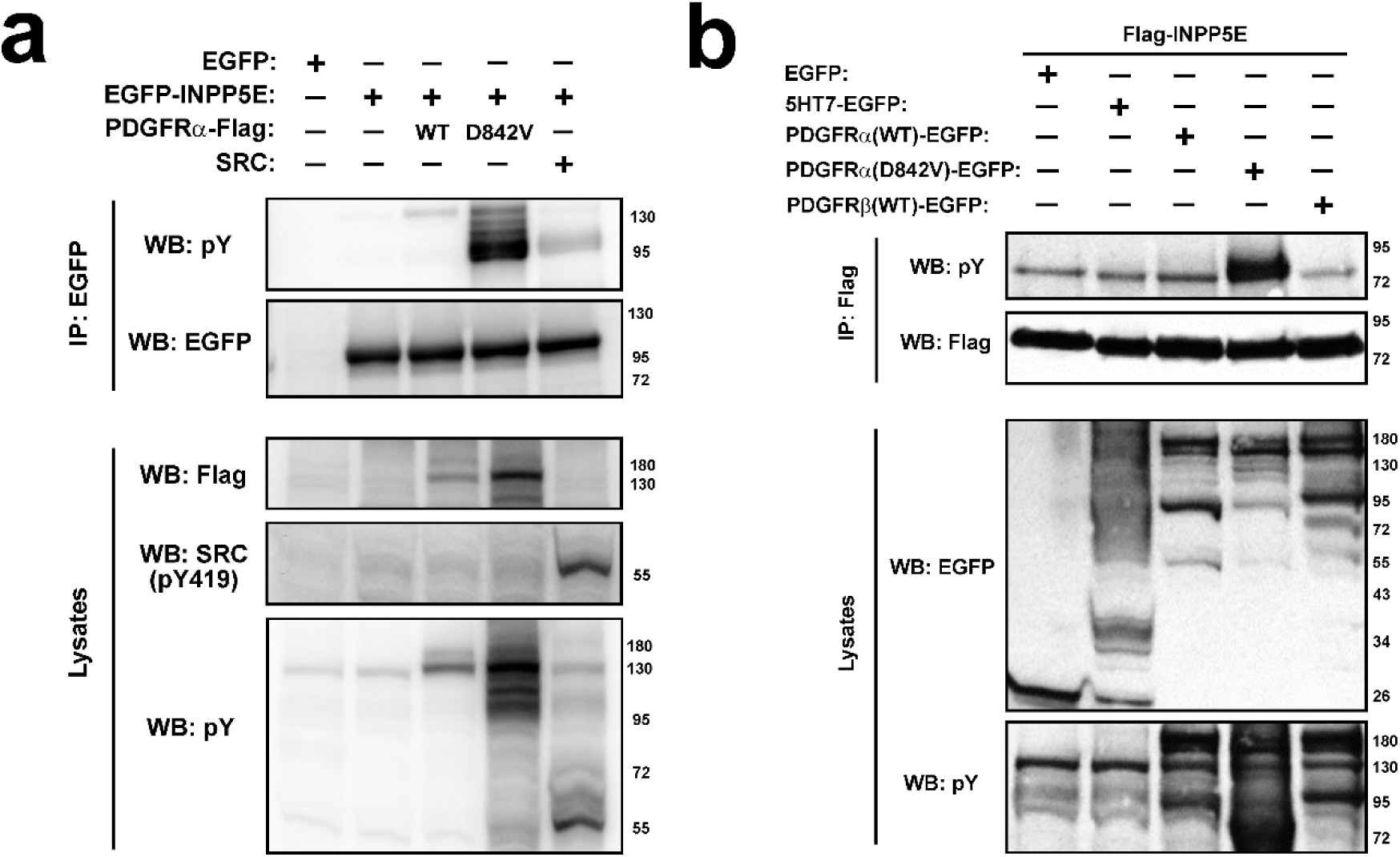
PDGFRα-D842V and SRC induce tyrosine phosphorylation of INPP5E. **(a)** Lysates from HEK293T cells transiently expressing the indicated proteins (top) were subjected to immunoprecipitation with anti-EGFP beads (IP: EGFP). Immunoprecipitates (IPs) and lysates were then analyzed by Western blot (WB) with antibodies against phosphotyrosine (pY), EGFP, Flag or active SRC (SRC pY419). **(b)** Similar experiment as in (a), except INPP5E was expressed with a Flag epitope and immunoprecipitated with anti-Flag beads (IP: Flag), while the other proteins were expressed as EGFP fusions. Molecular weight markers are shown on the right (kDa). Note how pY signal in (b) matches size of Flag-INPP5E (≈73 kDa), whereas in (a) it matches that of EGFP-INPP5E (≈100 kDa).

The above experiment suggested but did not prove that INPP5E undergoes tyrosine phosphorylation, as it could not rule out that the pY signal matching the EGFP-INPP5E band was due to phosphorylation of either the EGFP tag or a co-precipitating protein with the same electrophoretic mobility (≈100 kDa). We assessed these possibilities by switching tags: we now immunoprecipitated Flag-INPP5E (≈73 kDa) after coexpressing it with PDGFRα-EGFP (WT or D842V) (Fig.1b). When this was done, Flag-INPP5E displayed strong pY signal in presence of PDGFRα-D842V, but not with PDGFRα-WT, PDGFRβ-WT, the 5HT_7_ G protein-coupled receptor, or empty vector (Fig.1b) [49].

Since the robust D842V-induced pY band was observed with both EGFP-INPP5E (≈100 kDa) and Flag-INPP5E (≈73 kDa), these data strongly suggest that D842V promotes tyrosine phosphorylation of INPP5E itself.

We next asked whether D842V affects INPP5E catalytic activity. To test this, we performed phosphatase assays in anti-EGFP immunoprecipitates of HEK293T cells cotransfected with EGFP-INPP5E and PDGFRα-Flag (WT or D842V). As controls, we used cells transfected with EGFP-INPP5E-WT alone, or with EGFP-INPP5E-D477N, a catalytically inactive active site mutant [4]. As expected, INPP5E-WT but not INPP5E-D477N readily dephosphorylated PI(4,5)P_2_ and PIP_3_ in these assays. The activity of INPP5E-WT was very similar, if not the same, in presence of PDGFRα-D842V or WT, suggesting that INPP5E tyrosine phosphorylation does not affect its catalysis (Suppl. Fig. S1).

We then tested whether EGFP-INPP5E tyrosine phosphorylation is stimulated by orthovanadate, a broad specificity tyrosine phosphatase inhibitor [55]. This was indeed the case, with orthovanadate inducing strong pY signal not only on EGFP-INPP5E-WT, but also on its Y293F and Y614F non-phosphorylatable mutants, affecting exposed tyrosines in the catalytic domain (Y293) or in the FDRLYL motif key for ciliary targeting (Y614) (Suppl. Fig. S2) [4]. Hence, INPP5E tyrosine phosphorylation probably does not involve Y293 or Y614, but rather one or more of its remaining 14 tyrosines. Altogether, our data show that INPP5E undergoes tyrosine phosphorylation, which is strongly induced by PDGFRα-D842V.

### Effect of PDGFRα-D842V on the INPP5E interactome

To address how phosphotyrosine in INPP5E affects its interactions, we next characterized the interactome of Flag-INPP5E in HEK293T cells in presence of PDGFRα-EGFP (WT or D842V). As negative control, we also coexpressed PDGFRα-EGFP (D842V) with empty Flag vector. Mass spectrometric analysis of anti-Flag immunoprecipitates from these three samples showed robust and specific Flag-INPP5E immunoprecipitation (IP), as well as coprecipitation of several known INPP5E interactors, like PDE6D and 14-3-3 proteins (Table 1) [8, 56]. In addition, we identified multiple novel candidate INPP5E interactors, including SH3 domain proteins, protein phosphatase 2 subunits, mitochondrial proteins, chaperones, and many others (Table 1). Although most candidate interactors showed no signs of D842V-dependence, there were a few (e.g. SH3GL1, SNX9, STRAP) for which peptide count data suggested a possible increase, which will be addressed later. The full list of identified proteins is available as Suppl. Data File 1.

**Table 1.** Summary of INPP5E interactomics +/- PDGFRα-D842V. Left column includes the most specific hits, grouped in categories, from HEK293T proteomic study of Flag-INPP5E immunoprecipitates in presence of PDGFRα-EGFP (WT or D842V). Right three columns show peptide count in each of the samples, including negative control on the left. Full results of this experiment are available as Supplementary Data File 1.

| Identified protein | Peptide Count (IP: Flag) |  |  |
| --- | --- | --- | --- |
| | Flag empty vector + PDGFRα-EGFP (WT) | Flag-INPP5E + PDGFRα-EGFP (D842V) | Flag-INPP5E + PDGFRα-EGFP (WT) |
| **INPP5E** | 0 | 80 | 73 |
| **14-3-3 proteins:** |  |  |  |
| YWHAZ | 1 | 30 | 31 |
| YWHAH | 2 | 21 | 27 |
| YWHAQ | 2 | 26 | 30 |
| YWHAE | 1 | 35 | 39 |
| YWHAG | 2 | 24 | 24 |
| YWHAB | 3 | 26 | 25 |
| **Known interactors:** |  |  |  |
| PDE6D | 0 | 3 | 3 |
| USP9X | 0 | 3 | 3 |
| STUB1/CHIP | 0 | 10 | 6 |
| **SH3-containing:** |  |  |  |
| SH3GL1 | 0 | 4 | 1 |
| GRB2 | 0 | 2 | 3 |
| SNX9 | 0 | 2 | 0 |
| **PP2A subunits:** |  |  |  |
| PPP2R1A | 1 | 17 | 11 |
| PPP2R2A | 0 | 13 | 5 |
| PPP2CB | 0 | 4 | 3 |
| **Mitochondrial:** |  |  |  |
| SLC25A13 | 0 | 9 | 8 |
| SLC25A12 | 0 | 3 | 4 |
| SLC25A11 | 1 | 9 | 11 |
| TIMM50 | 0 | 6 | 7 |
| BCKDK | 0 | 7 | 11 |
| **Chaperones:** |  |  |  |
| DNAJC10 | 0 | 5 | 5 |
| DNAJA1 | 1 | 10 | 9 |
| DNAJA3 | 0 | 3 | 3 |
| HSPH1 | 0 | 14 | 4 |
| HSPA4 | 0 | 7 | 4 |
| **Others:** |  |  |  |
| STRAP | 0 | 13 | 6 |
| PKM | 1 | 10 | 5 |
| TXNDC5 | 0 | 10 | 4 |
| EEF2 | 2 | 12 | 8 |
| EIF4A1 | 0 | 8 | 1 |
| PFN2 | 0 | 3 | 4 |

### Serine 85 of INPP5E is essential for interaction with 14-3-3ζ

Six of the seven human 14-3-3 isoforms were the most abundant INPP5E-coprecipitating proteins in our interactome (Table 1), consistent with a previous study [8]. 14-3-3 proteins typically bind to specific phosphoserine (pS) or phosphothreonine (pT)-containing motifs on their targets, thereby regulating their functions [30]. We therefore looked for potential 14-3-3-binding sites on INPP5E. According to the Eukaryotic Linear Motif (ELM) resource, INPP5E contains six canonical 14-3-3-binding motifs (Fig. 2a) [57]. We then used the PhosphoSitePlus database to establish which of these INPP5E’s putative 14-3-3-binding motifs are known to undergo phosphorylation [58]. Interestingly, the first of these motifs (82-RAL<u>S</u>LD-87) contains S85, one of the three most consistently phosphorylated residues in INPP5E (the others being S99 and S241, which are not part of such motifs) (Fig. 2a). Hence, S85 is a good candidate to regulate 14-3-3 protein binding.

**Figure 2.**
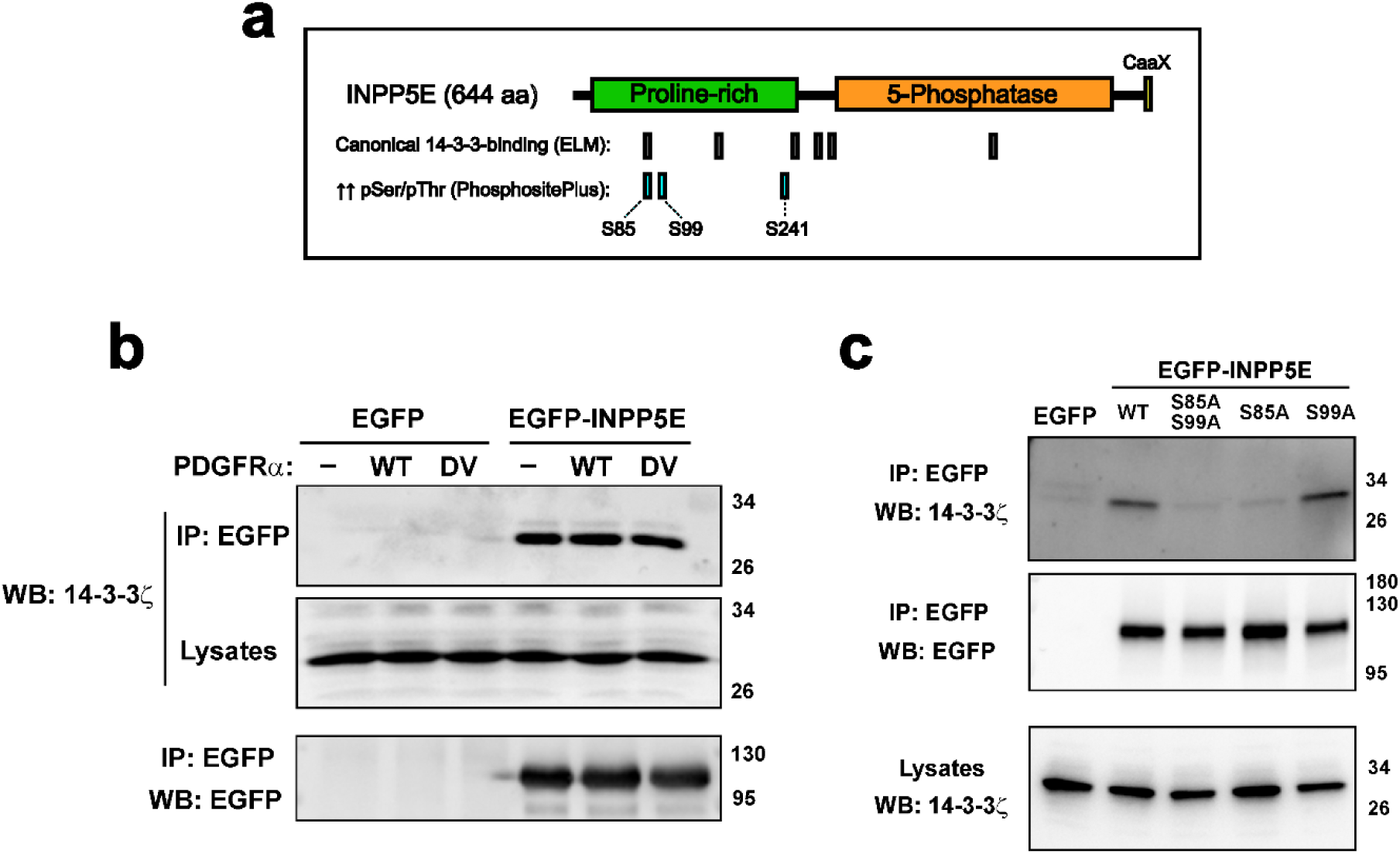
INPP5E interacts with 14-3-3ζ in a serine 85-dependent manner. **(a)** Schematic illustrating INPP5E’s domain structure. Also shown are its six canonical 14-3-3-binding sites (according to ELM program) and its three most phosphorylated residues (according to PhosphoSitePlus database). **(b)** Co-immunoprecipitation (co-IP) experiment in HEK293T cells shows EGFP-INPP5E interacts with endogenous 14-3-3ζ regardless of PDGFRα expression in its wild type (WT) or D842V mutant (DV) form. **(c)** HEK293T Co-IP of endogenous 14-3-3ζ by EGFP-INPP5E is affected by S85 but not S99 mutation to alanine. Molecular weight markers shown on the right.

We next used a specific anti-14-3-3ζ antibody to confirm its interaction with INPP5E. Indeed, endogenous 14-3-3ζ robustly coprecipitated with EGFP-INPP5E in a PDGFRα-D842V-independent manner, in line with our proteomic data (Fig. 2b, Table 1). We then checked for dependence on S85 and S99, which we mutated to non-phosphorylatable alanines. 14-3-3ζ readily interacted with the S99A mutant of EGFP-INPP5E, but not with the S85A mutant, or with the double S85A+S99A mutant (Fig. 2c). Hence, we conclude that S85 but not S99 is essential for INPP5E binding to 14-3-3ζ.

### INPP5E interacts with SIN1, USP9X, STRAP and CAMSAP2

By looking at our list of putative INPP5E interactors, as well as those of previous studies [8, 56, 59], we then selected a few candidates for further study, with the main selection criterion being the existence of known connections to INPP5E-related structures or processes, such as cilia, microtubules, phosphoinositides or growth factor signaling. Aside from SH3 domain-containing proteins, dealt with in the next section, the selected proteins were: USP9X (a deubiquitylase involved in centrosome and cilia biology), STRAP (a regulator of TGFβ and phosphoinositide signaling), SIN1 (a phosphoinositide-binding mTORC2 complex subunit), CAMSAP2 (a microtubule nucleator), PKM2 (a phosphotyrosine-binding glycolytic enzyme), CHIP (an E3 ubiquitin ligase regulating cilia resorption), and CEP170 (another cilia resorption regulator) [32–35, 60–68].

By using antibodies against these proteins, we first assessed which ones coprecipitated with EGFP-INPP5E in HEK293T cells, and whether the association was affected by PDGFRα-WT or D842V. Endogenous USP9X, STRAP, SIN1 and CAMSAP2 all specifically coprecipitated with EGFP-INPP5E, regardless of PDGFRα status (Fig. 3a-c), whereas no coprecipitation was observed for CEP170, PKM2 and CHIP (not shown).

**Figure 3.**
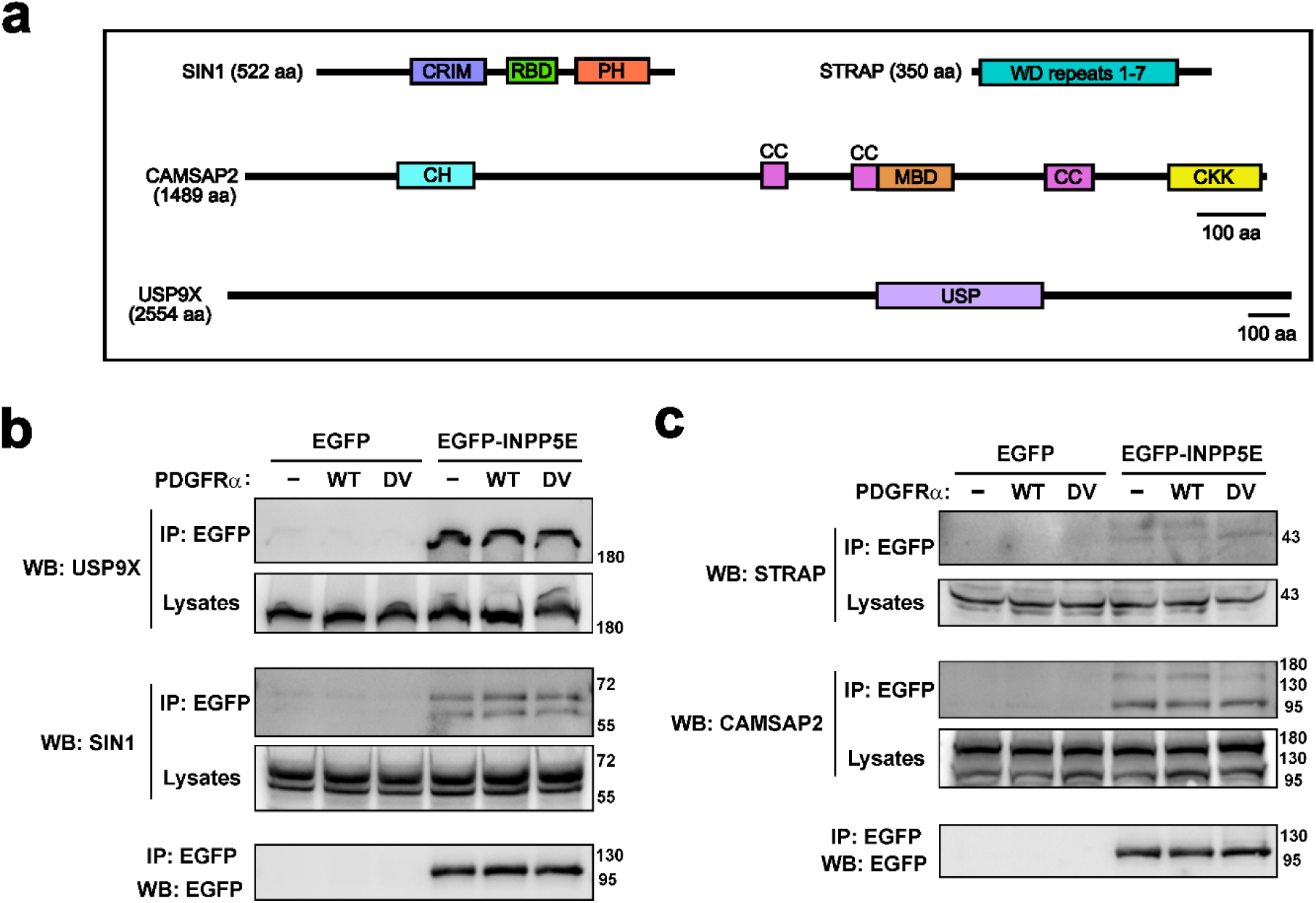
INPP5E interacts with USP9X, SIN1, STRAP and CAMSAP2. **(a)** Schematic of SIN1, STRAP, CAMSAP2 and USP9X showing their conserved domains according to UniProt database. CRIM: conserved region in the middle; RBD: Ras-binding domain; PH: Pleckstrin-homology; CH: Calponin homology; CC: Coiled coils; MBD: microtubule-binding domain; CKK: microtubule minus end recognition; USP: ubiquitin-specific protease. Scale bars are shown (USP9X drawn at smaller scale). **(b)** HEK293T co-IP of endogenous USP9X and SIN1by EGFP-INPP5E, in presence or not of PDGFRα-Flag (WT or D842V mutant (DV)). **(c)** Co-IP as in (b) of endogenous STRAP and CAMSAP2. Molecular weight markers on the right.

For the confirmed interactors, we also checked whether protein levels or subcellular localization were affected in INPP5E-KO versus WT hTERT-RPE1 cells [4]. Western blot of USP9X, STRAP, SIN1 and CAMSAP2 revealed no differences, and the same was true for 14-3-3ζ, and for SH3 proteins SH3GL1 and SNX9 (see next section) (Suppl. Fig. S3). Immunofluorescence for all these proteins in these same cells also revealed no differences (not shown). Altogether, our data suggest that INPP5E interacts with USP9X, STRAP, SIN1 and CAMSAP2, without affecting their protein levels or localization.

### INPP5E interacts with SH3 proteins SNX9, SH3GL1 and GRB2

INPP5E contains three putative SH3-binding motifs in its proline-rich region (Fig. 4a) [6]. Consistently, several SH3 domain proteins appeared as candidate INPP5E interactors in our proteomic experiment, including GRB2, SNX9 and SH3GL1 (also known as Endophilin A2) (Fig. 4b). EYFP-tagged versions of these three proteins all robustly coprecipitated Flag-INPP5E in HEK293T cells, confirming the interactions (Fig. 4c). Moreover, for SNX9 and SH3GL1, but not GRB2, the interaction appeared stronger in presence of PDGFRα-D842V, as compared to PDGFRα-WT (Fig. 4d). Thus, SNX9 and SH3GL1 may warrant further study as candidate effectors of INPP5E tyrosine phosphorylation.

**Figure 4.**
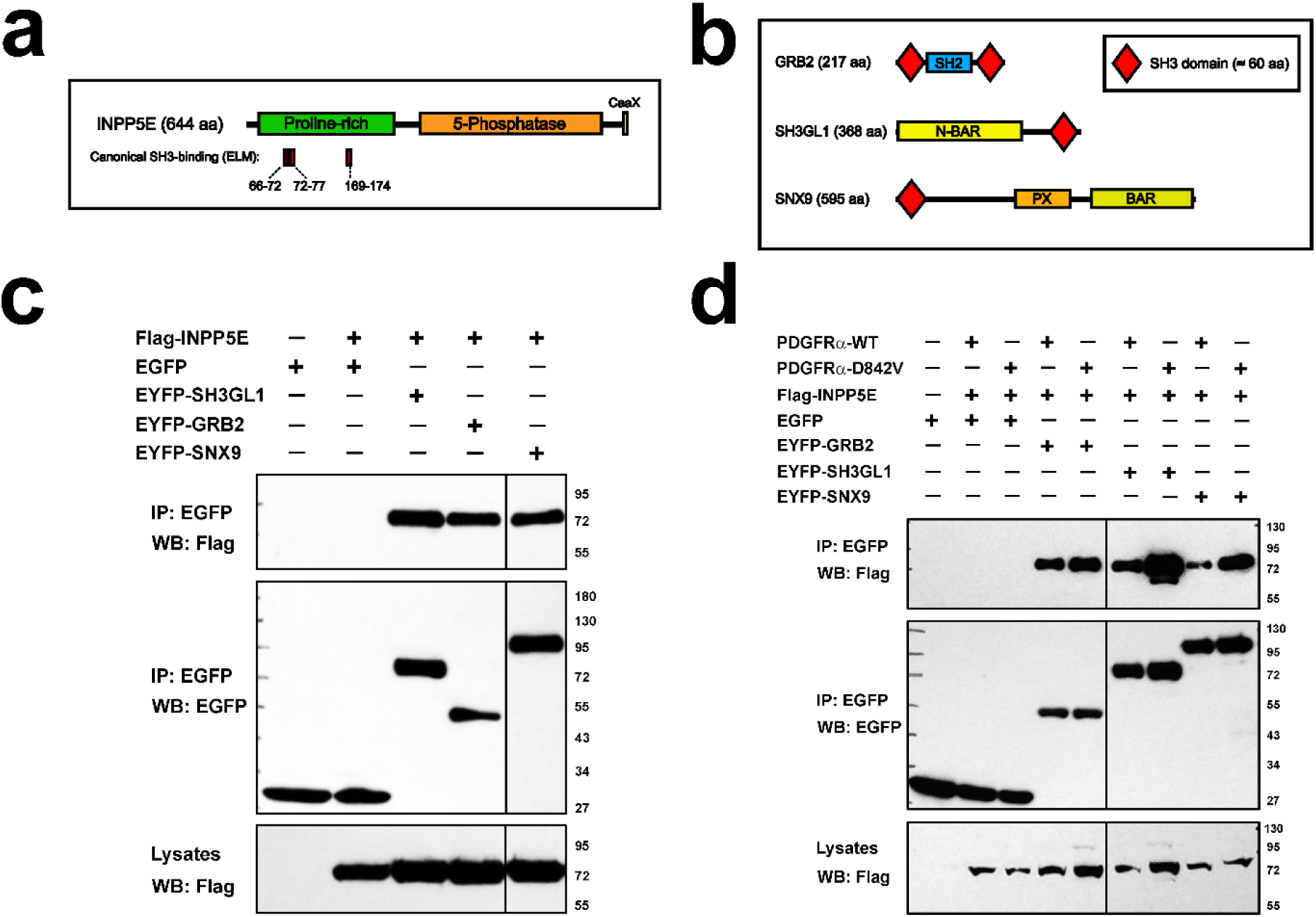
INPP5E interacts with SNX9, SH3GL1 and GRB2. **(a)** Diagram of INPP5E showing its three canonical SH3-binding sites within its proline-rich region (based on ELM). **(b)** Schematic of GRB2, SH3GL1 and SNX9 showing their SH3 and other domains (based on UniProt). **(c)** HEK293T co-IP showing association of Flag-INPP5E with EYFP-tagged SH3GL1, GRB2 and SNX9. **(d)** Co-IP experiment as in (c) to assess effect of PDGFRα-Flag (WT or D842V) on the interactions from (c). Molecular weight markers on the right.

### INPP5E interacts with Joubert syndrome proteins NPHP1 and AHI1

NPHP1 and AHI1 (also known as Jouberin) are two JBTS-causative transition zone proteins required for INPP5E ciliary targeting, and both contain SH3 domains (Fig. 5a) [7, 8, 69–73]. We therefore wondered whether they interact with INPP5E, and whether such interactions depend on any of the four ciliary localization signals (CLS1-4) we previously characterized in INPP5E [4].

**Figure 5.**
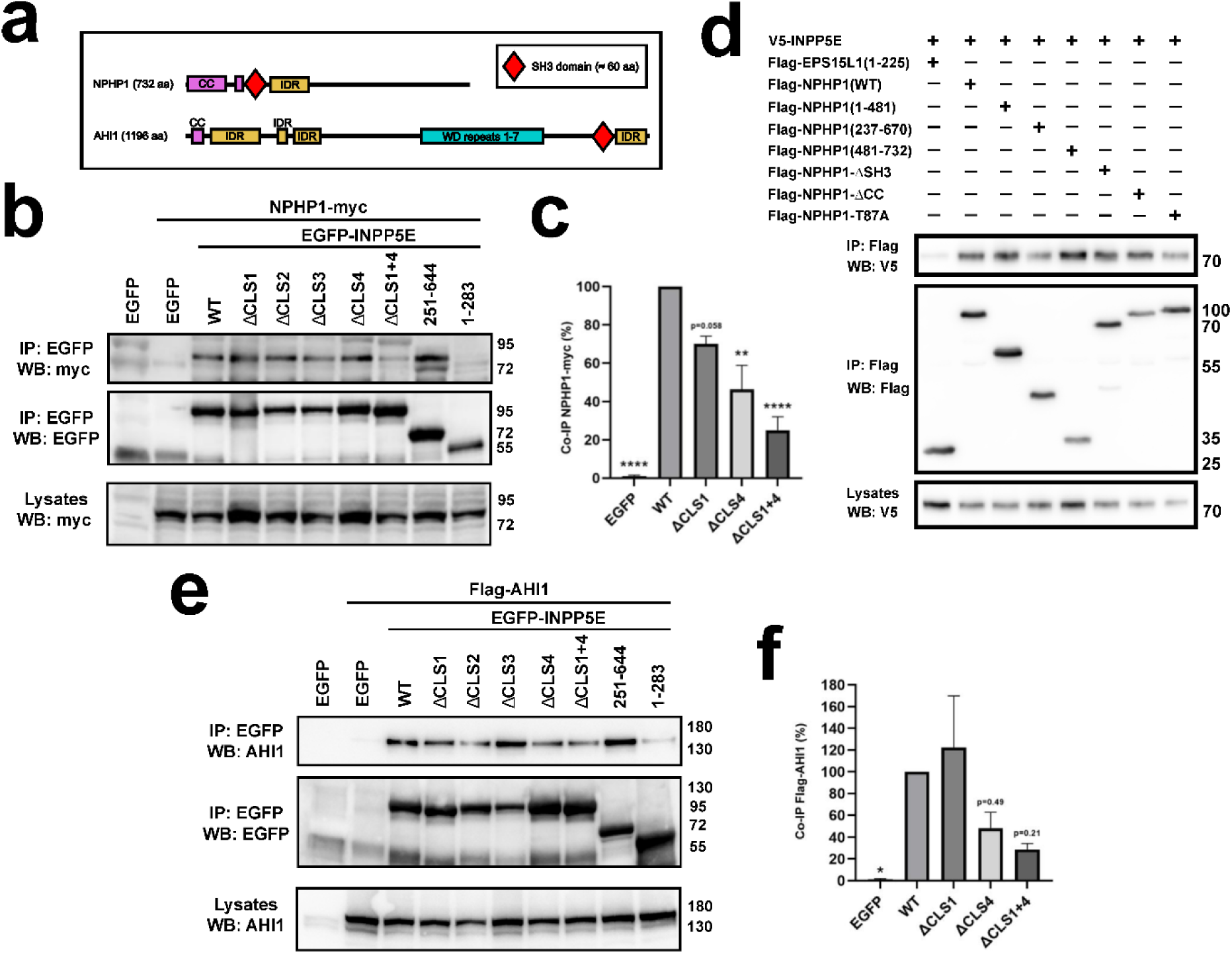
INPP5E interacts with NPHP1 and AHI1/Jouberin. **(a)** Diagram of NPHP1 and AHI1 displaying their SH3 domains, coiled coils (CC) and intrinsically disordered regiones (IDRs) (based on UniProt). **(b)** HEK293T co-IP of NPHP1-myc by EGFP-INPP5E WT and its previously reported ciliary localization signal (CLS) and deletion mutants. **(c)** Quantitation NPHP1-myc co-IP with EGFP-INPP5E WT and indicated mutants. Data are mean ± SEM of n=3 independent experiments, including the one in (b), and were analyzed by one-way ANOVA followed by Tukey tests (p<0.01: **; p<0.0001: ****). **(d)** HEK293T co-IP of V5-INPP5E by Flag-NPHP1 WT and its previously reported mutants, including deletions and the T87A point mutant, which deletes a PLK1 phosphorylation site. Flag-EPS15L1(1-225) was used as negative control. **(e)** Co-IP as in (c) of Flag-AHI1 by EGFP-INPP5E and its mutants. **(f)** Quantitation as in (d) of Flag-AHI1 co-IP (p<0.05: *).

In HEK293T, NPHP1-myc readily coprecipitated with EGFP-INPP5E-WT, and with the individual CLS mutants from the above study: ΔCLS1, ΔCLS2, ΔCLS3 and ΔCLS4 (Fig. 5b). However, upon quantitation of several experiments, it became clear that the interaction is reduced with ΔCLS4, and even more so in the double ΔCLS1+4 mutant (Fig. 5c). Somewhat surprisingly, residues 251-644 of INPP5E (encompassing catalytic domain and all CLSs, but lacking all SH3-binding motifs) was sufficient for the NPHP1-INPP5E interaction, whereas residues 1-283 (including all such motifs) were not (Fig. 5b).

We then explored how different NPHP1 regions affect its INPP5E interaction. Flag-NPHP1 (732 aa) specifically coprecipitated V5-tagged INPP5E, and so did each of six previously described Flag-NPHP1 mutants [50], including three deletions together encompassing the full protein (1-481, 237-670, 481-732), a mutant lacking the coiled coil (ΔCC), another lacking the SH3 domain (ΔSH3), and the T87A point mutant lacking a PLK1 phosphorylation site (Fig. 5d). Flag-EPS15L1(1-225) was used as negative control and showed no interaction [50]. Since the NPHP1-INPP5E interaction was not abolished by any of these NPHP1 mutants, we hypothesize that INPP5E concomitantly binds to more than one NPHP1 region, a possibility awaiting further exploration.

Like NPHP1-myc, Flag-AHI1 also coprecipitated robustly with EGFP-INPP5E, with the interaction showing a downward trend in mutants lacking CLS4 (i.e. the CaaX box) (Fig. 5e-f). Also like NPHP1, INPP5E residues 251-644 sufficed for the AHI1 interaction, whereas only weak binding was seen with residues 1-283 (Fig. 5e).

### INPP5E downregulates PDGF-AA signaling and PDGFRα receptor levels

PDGFRα-D842V’s effect on INPP5E’s tyrosine phosphorylation suggests a possible role for INPP5E in PDGF signaling. To test this, we studied how PDGF-AA treatment for 0, 10, 30 and 60 minutes regulates AKT phosphorylation in *Inpp5e*-null (Inpp5e-KO) and heterozygous littermate control (Inpp5e-Het) mouse embryonic fibroblasts (MEFs) [13]. We looked at the two main phosphorylations of AKT, both of which depend on PIP_3_: S473 phosphorylation by mTORC2 (p-S473), and T308 phosphorylation by PDK1 (p-T308) [35].

In control MEFs, PDGF-AA moderately increased p-S473 and p-T308 at 10 min of treatment, after which both phosphorylations went down. In Inpp5e-KO MEFs, by contrast, induction of p-S473 and p-T308 was much more robust, with maximal levels at 10 min significantly higher than in control (Fig. 6a-c). As additional control, we also treated cells for 10 min with fetal bovine serum (FBS), which induced AKT phosphorylations equally robustly in both cell types (Fig. 6a). Hence, INPP5E downregulates AKT phosphorylation in response to PDGF-AA but not FBS.

**Figure 6.**
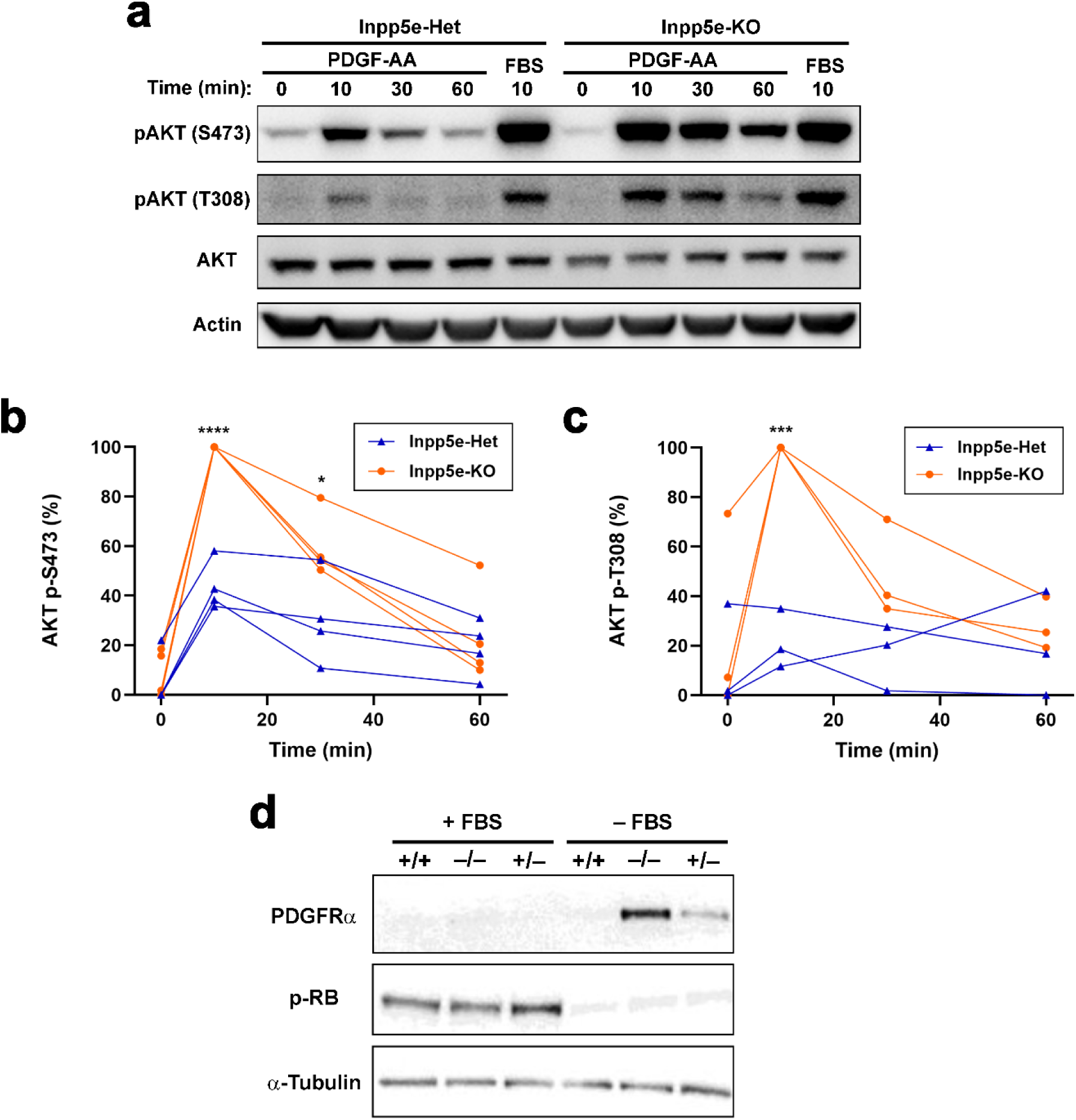
INPP5E downregulates PDGF signaling. **(a)** *Inpp5e*^+/‒^ and *Inpp5e*^‒/‒^ MEFs were stimulated with PDGF-AA or FBS for the indicated times and cell lysates analyzed by Western blot with the indicated antibodies. Actin was used as loading control. **(b-c)** Quantitations of normalized pAKT/AKT ratios from n=4 (S473) or n=3 (T308) independent experiments, including the one in (a). **(d)** *Inpp5e*^+/+^, *Inpp5e*^+/‒^ and *Inpp5e*^‒/‒^ MEFs cultured with or without FBS were analyzed by Western blot with the indicated antibodies. Tubulin was used as loading control and phospho-retinoblastoma (pRb) as cell cycle marker. Note INPP5E-dependence of PDGFRα receptor levels in serum-starved MEFs.

To better understand INPP5E’s effects on PDGF signaling, we also looked at the protein levels of the PDGF-AA receptor, PDGFRα, in the above Inpp5e-Het and Inpp5e-KO littermate MEFs, as well as in non-littermate Inpp5e-WT MEFs. When grown in FBS, all these MEF lines were actively going through the cell cycle (as seen by high phospho-retinoblastoma levels, p-Rb). Under these conditions, PDGFRα levels were virtually undetectable, as one might expect given high growth factor levels in the medium. In contrast, in serum-starved MEFs, we observed an inverse correlation between INPP5E dose and PDGFRα levels: highest in Inpp5e-KO, intermediate in Inpp5e-Het, and lowest in WT (Fig. 6d). This may be part of the reason why serum-starved Inpp5e-KO MEFs responded more strongly to PDGF-AA (Fig. 6a-c).

To explore growth factor specificity, we also treated MEFs with insulin instead of PDGF. Unlike PDGF, insulin induction of p-S473 in AKT occurred equally intensely in Inpp5e-Het and KO MEFs (Suppl. Fig. S4). Lastly, we also analyzed AKT phosphorylation in INPP5E-KO and WT hTERT-RPE1 cells, which, unlike MEFs, are human epithelial cells [4]. For this, we stimulated these cells for 10 minutes with PDGF-AA, IGF-1, EGF or FBS. Consistent with almost undetectable expression of PDGFRα in this cell type [51], PDGF-AA failed to elicit AKT phosphorylation. In contrast, IGF-1, EGF and FBS all induced robust AKT phosphorylation of S473 and T308, but did so equally in INPP5E-KO and WT (Suppl. Fig. S5). Therefore, INPP5E downregulation of PDGF-induced AKT phosphorylation in MEFs is both ligand- and cell type-specific.

### INPP5E promotes TGFβ signaling

We next tested whether INPP5E affects TGFβ signaling, another cilia-dependent growth factor signaling pathway [2, 41–44]. For this, we treated Inpp5e-Het and KO MEFs with TGFβ for 0, 10, 30, 60, 90 and 180 min and looked at induction of SMAD2 and ERK phosphorylations. In Inpp5e-Het MEFs, TGFβ treatment led to accumulation of phospho-SMAD2 (p-SMAD2) during the first 30 min, after which its levels stayed high for an hour (30-90 min) and then went down at 180 min. By contrast, p-SMAD2 induction was strongly attenuated in Inpp5e-KO MEFs (Fig. 7a-b). As for phospho-ERK (p-ERK), its levels reached a maximum at 180 min in Inpp5e-Het MEFs, and again this increase was significantly less pronounced in Inpp5e-KO MEFs (Fig. 7a,c). Thus, INPP5E is a positive regulator of TGFβ signaling in fibroblasts.

**Figure 7.**
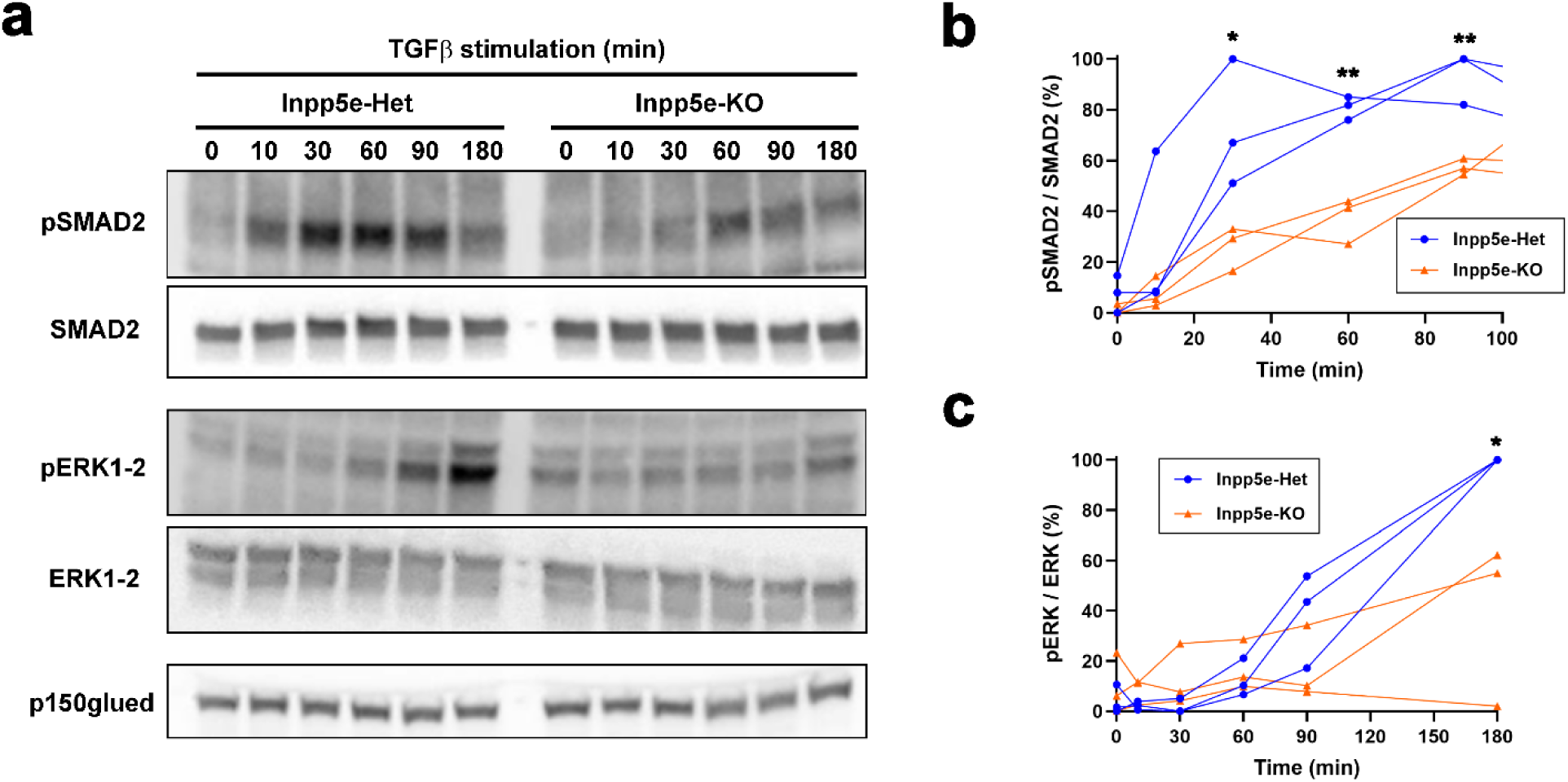
INPP5E promotes TGFβ signaling. **(a)** *Inpp5e*^+/‒^ and *Inpp5e*^‒/‒^ MEFs were stimulated with TGFβ for the indicated times and cell lysates analyzed by Western blot with the indicated antibodies. p150glued was used as loading control. **(b)** Quantitation of normalized phospho-SMAD2 to total SMAD2 ratios from n=3 independent experiments, including the one in (a). **(c)** Quantitation of normalized phospho-ERK1-2 to total ERK1-2 ratios from n=3 independent experiments, including the one in (a). Data in b-c were analyzed by two-tailed unpaired t-tests (p<0.05: *; p<0.01: **).

## Discussion

Here, we studied the role of INPP5E in growth factor signaling. First, we discovered that INPP5E is strongly tyrosine-phosphorylated in presence of the constitutively active PDGFRα-D842V, and less intensely by wild type SRC tyrosine kinase (Fig.1). D842V-induced tyrosine phosphorylation did not affect INPP5E enzyme activity (Fig. S1), so we explored its effects on INPP5E’s interactome. Although most INPP5E interactors were D842V-independent, our proteomic and co-IP experiments suggested that INPP5E’s interaction with two SH3 proteins, SNX9 and SH3GL1, is stronger with D842V (Table 1, Fig. 4).

All this points to a role for INPP5E in receptor tyrosine kinase (RTK) signaling. Accordingly, we found that INPP5E-KO MEFs respond more strongly to PDGF-AA and contain higher levels of its receptor PDGFRα (Fig. 6). Interestingly, SH3GL1 promotes endocytosis of some RTKs, so it might conceivably be involved in INPP5E-dependent PDGFRα downregulation [74, 75]. As for SNX9, it binds phosphoinositides, modulates F-actin, and localizes to sites of ciliary vesicle scission, an INPP5E-regulated process [28, 76, 77]. Thus, the SNX9-INPP5E interaction may hold important clues about how growth factors trigger ciliary vesicle release.

Our data also suggest that INPP5E is regulated by serine phosphorylation. Consistent with previous work, we found that INPP5E strongly interacts with 14-3-3 proteins (Table 1) [8, 56]. Of the seven human 14-3-3 proteins (β, γ, ε, ζ, η, θ, σ), INPP5E interacted with all except σ, and all interactions were pY-independent (Table 1; Fig. 2b). 14-3-3 proteins bind over a thousand mammalian proteins, and do so via specific pS/pT-containing phosphopeptides [30]. INPP5E has six such canonical 14-3-3-binding sites, but only the one containing S85 is known to be phosphorylated (Fig. 2a). Consistently, the non-phosphorylatable S85A mutant failed to bind 14-3-3ζ (Fig. 2c). Since 14-3-3 binding typically regulates the activity or interactions of their target proteins, this suggests S85 phosphorylation status as an additional mode of INPP5E regulation. As of this writing, PhosphoSitePlus points to 24 different studies identifying S85 phosphorylation in INPP5E, suggesting that this serine is normally phosphorylated [58]. For instance, a phosphoproteomic study of short-term Hedgehog signaling found similar S85 phosphorylation levels under basal and stimulated conditions [78]. If S85 is indeed phosphorylated by default, then its regulation might occur via dephosphorylation, perhaps via kinase inhibition or localized activation of PP2A, a protein phosphatase whose subunits featured prominently in our interactome (Table 1). Consistent with this hypothesis, PP2A localizes to the ciliary base and regulates ciliary signaling via dephosphorylation of substrates such as KIF7 [39, 79, 80]. Future studies should clarify the upstream regulation and downstream effects of S85 phosphorylation and 14-3-3 binding to INPP5E.

Besides SNX9 and SH3GL1, we identified three other SH3 proteins as INPP5E interactors: GRB2, AHI1 and NPHP1 (Fig. 4–5). GRB2 is an adaptor that could conceivably target INPP5E to activated RTK complexes [81], whereas AHI1 and NPHP1 are JBTS-associated transition zone proteins needed for INPP5E ciliary targeting [7, 8, 69, 71–73]. Accordingly, we found that AHI1 and NPHP1 interact very poorly with ΔCLS1+4, an INPP5E mutant that fails to accumulate in cilia (Fig. 5) [4]. NPHP1 and AHI1 are also involved in growth factor signaling via tyrosine kinases, another possible connection to INPP5E [82–85].

Our co-IPs also confirmed INPP5E interactions with CAMSAP2, SIN1, USP9X and STRAP (Fig. 3) [8, 56]. CAMSAP2 is a microtubule minus end-stabilizing protein that promotes nucleation of microtubules, including those of motile cilia’s central pair [68]. SIN1 is the PIP_3_-binding subunit of mTORC2, the kinase mediating AKT phosphorylation at S473, which we found upregulated in PDGF-treated INPP5E-KO MEFs (Fig. 6) [35]. USP9X is a deubiquitinating enzyme that regulates centriolar satellites and ciliogenesis [60–63]. STRAP is both a repressor of TGFβ signaling and an activator of the PIP_3_-dependent kinase PDK1, responsible for AKT phosphorylation at T308 [32–34]. Hence, STRAP activation could potentially explain the TGFβ and PDGF responses we observed in INPP5E-KO MEFs (Fig. 6–7). This, however, remains to be determined, as is the functional relationship between INPP5E and all the interactors described herein. Altogether, our results shed new light into INPP5E’s involvement in growth factor signaling, and will help guide future studies on the molecular bases of ciliopathies.

## Acknowledgements

We thank Drs. Jorge Martín-Pérez, José M. Rojas and Takanari Inoue for reagents. This work was supported by grants from Spanish Ministerio de Ciencia e Innovación (MCIN) / Agencia Estatal de Investigación (AEI), and the European Regional Development Fund (FEDER) to F.R.G. (PID2023-149472NB-I00 from MCIN/AEI/10.13039/501100011033/FEDER, UE) and W.L. (PID2022-136654OB-I00 from MCIN/AEI/10.13039/501100011033/FEDER, UE). L.B.P. acknowledges funding from the Novo Nordisk Foundation (grants NNF18SA0032928 and NNF22OC0080406), the Independent Research Fund Denmark (grant 2032-00115B), the TheRaCil consortium funded by the European Union (Horizon-health-2022-disease-06-two stage, grant 101080717), and the Carlsberg Foundation (grant CF22-0670). J.L.B., F.I. and M.C. were funded by Agencia Nacional de Investigación e Innovación (ANII) and Programa para el Desarrollo de las Ciencias Básicas, Uruguay. F.I. was also funded by Comisión Sectorial de Investigación Científica (CSIC), Universidad de la República, Uruguay. B.S. was funded by the German Research Foundation (DFG; FOR5547; SCHE1562/12-1).

## Supplementary Figures

**Supplementary Figure S1.**
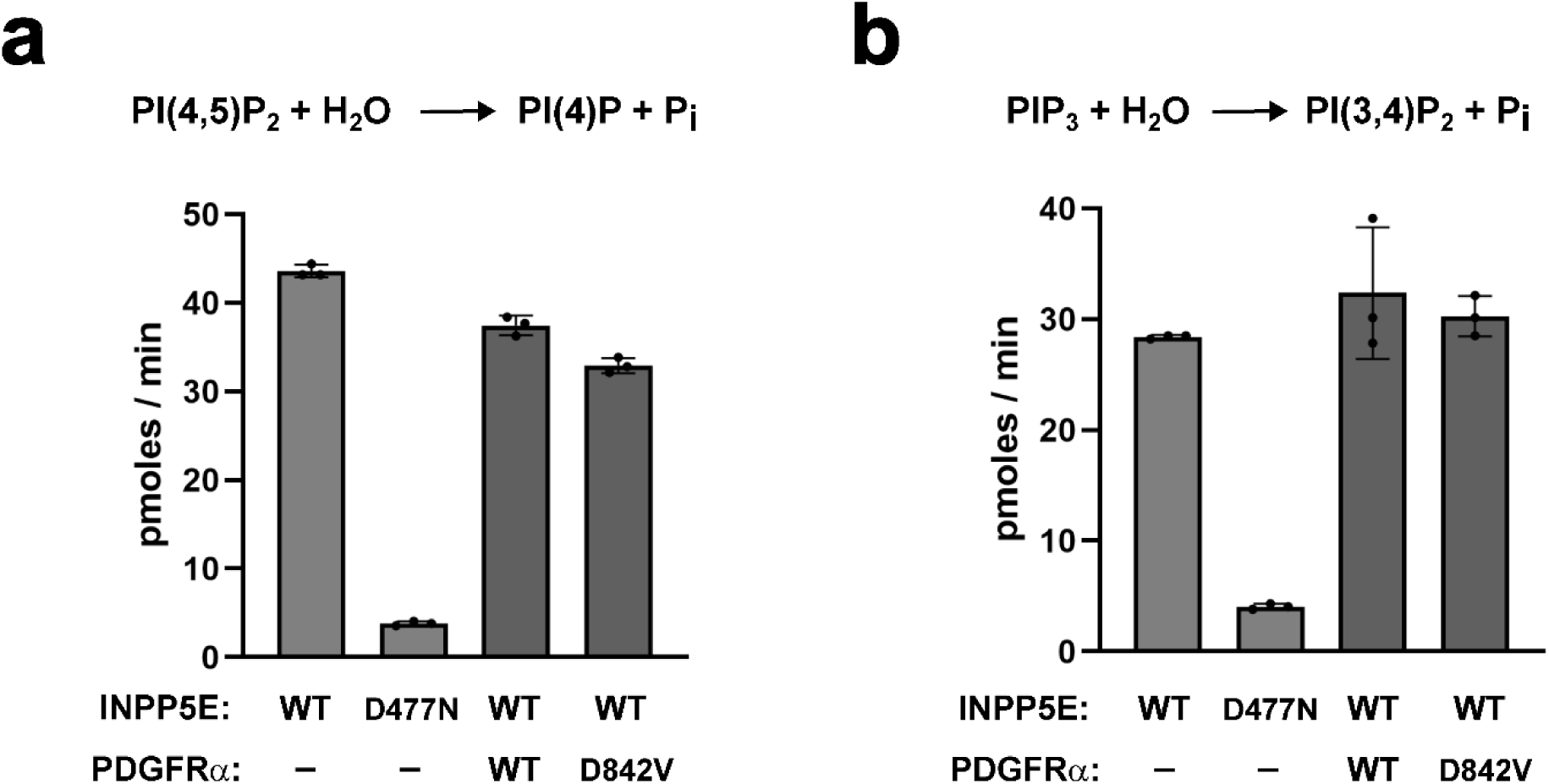
PDGFRα-D842V has no effect on INPP5E phosphatase activity. **(a)** PI(4,5)P_2_ phosphatase activity assay was performed on anti-EGFP immunoprecipitates from HEK293T cell lysates cotransfected with EGFP-INPP5E and PDGFRα-Flag (WT or D842V), as indicated. EGFP-INPP5E WT and D477N (catalytically inactive mutant) were used as positive and negative controls, respectively. Data are mean±SD of n=3 technical replicates from a single experiment. **(b)** Same as in (a) but using PIP_3_ as substrate for the phosphatase activity assay. The INPP5E-catalyzed 5-phosphatase reactions are shown above for both substrates.

**Supplementary Figure S2.**
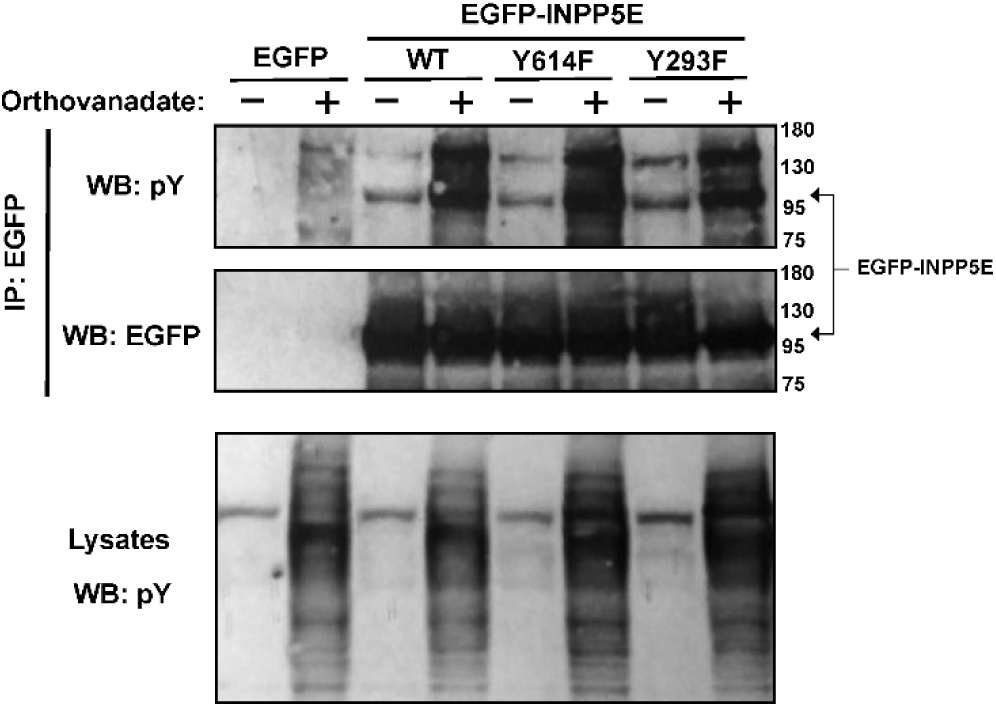
Orthovanadate increases tyrosine phosphorylation of INPP5E. **(a)** Lysates from HEK293T cells transiently expressing the indicated EGFP fusion proteins (top) and treated or not with orthovanadate, a tyrosine phosphatase inhibitor, were immunoprecipitated with anti-EGFP beads (IP: EGFP) and analyzed by Western blot (WB) with anti-phosphotyrosine (pY) and anti-EGFP antibodies, as indicated. Molecular weight markers on the right (kDa).

**Supplementary Figure S3.**
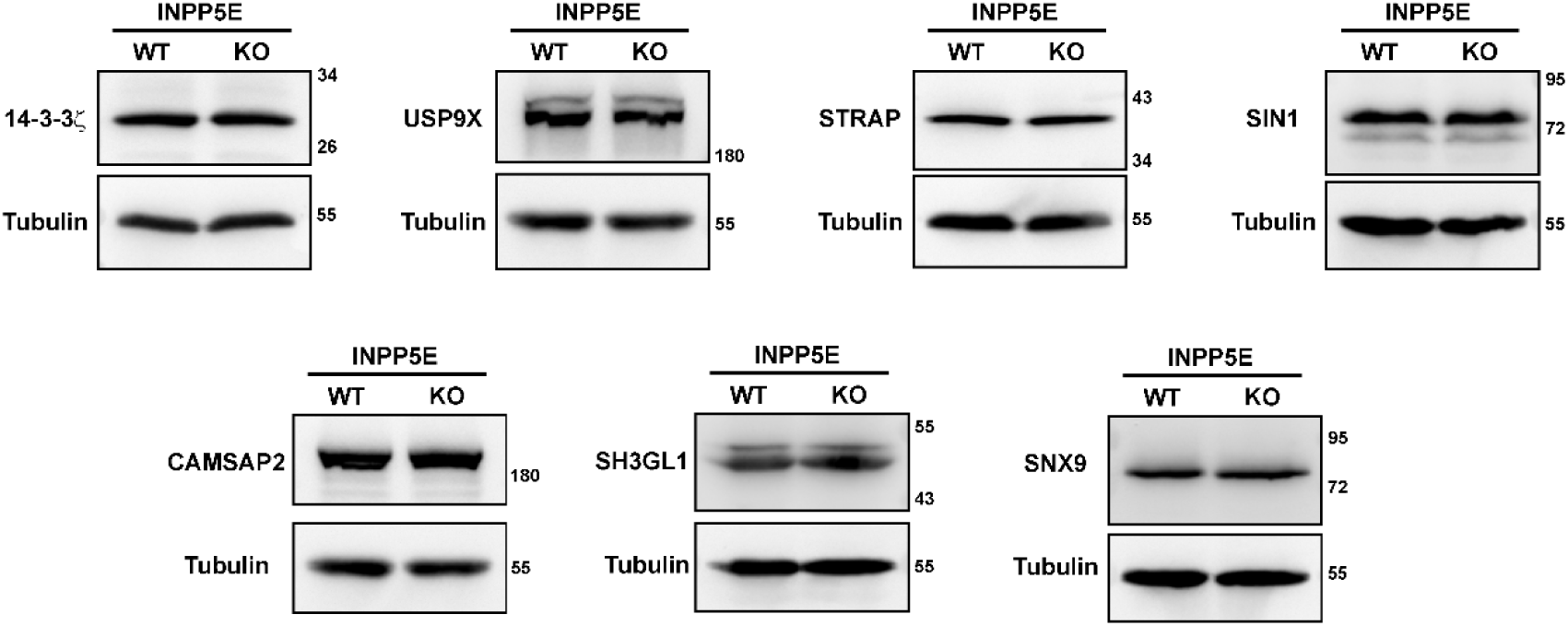
Protein levels of INPP5E interactors in INPP5E-KO cells. Lysates from INPP5E-WT and KO hTERT-RPE1 cells were analyzed by Western blot with the indicated antibodies against confirmed INPP5E interactors. Tubulin was used as loading control. Molecular weight markers on the right.

**Supplementary Figure S4.**
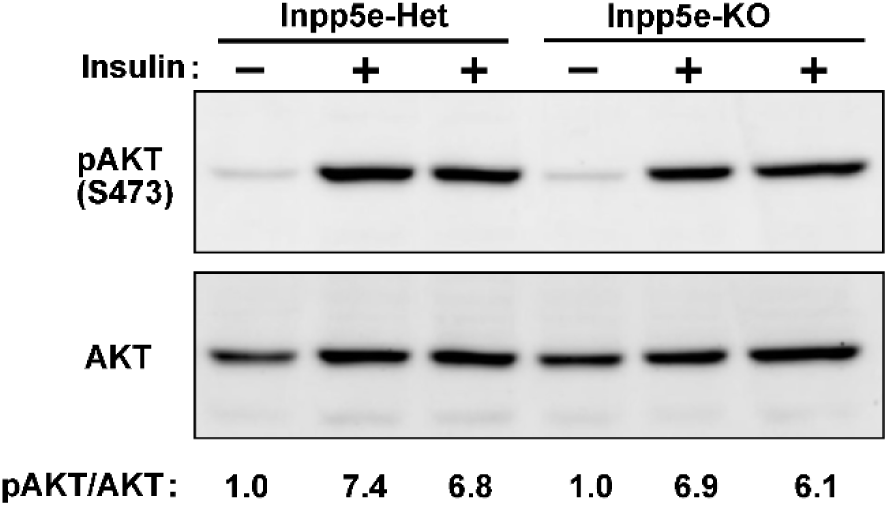
INPP5E has no effect on insulin-induced AKT phosphorylation in MEFs. *Inpp5e*^+/‒^ and *Inpp5e*^‒/‒^ MEFs were stimulated with insulin as indicated and AKT phosphorylation at S473 was analyzed by Western blot. The pAKT/AKT ratios for each sample, normalized to untreated heterozygous MEFs, are shown at the bottom.

**Supplementary Figure S5.**
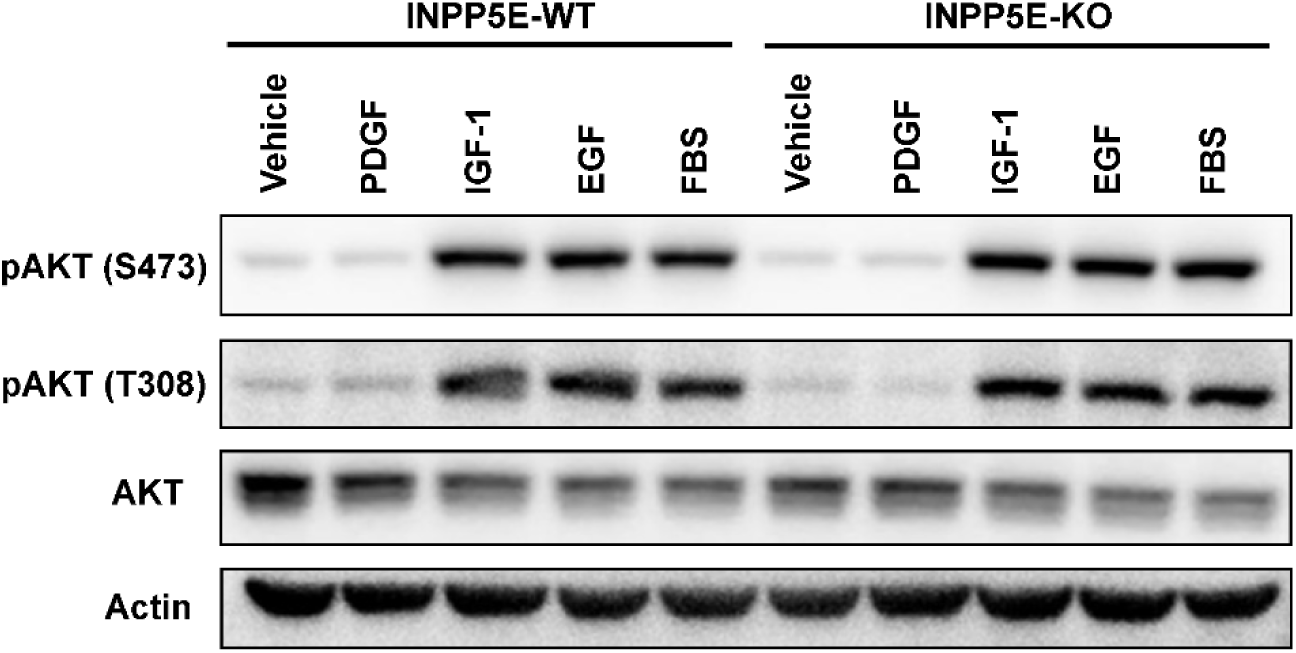
INPP5E has no effect on IGF- and EGF-induced AKT phosphorylation in hTERT-RPE1 cells. *INPP5E*^+/+^ and *INPP5E*^‒/‒^ hTERT-RPE1 cells were stimulated for 10 minutes with the indicated ligands and AKT phosphorylation at S473 and T308 was analyzed by Western blot, using total AKT and actin as controls.

